# Soluble epoxide hydrolase in Alzheimer’s disease drives neurovascular dysfunction

**DOI:** 10.64898/2025.12.09.692941

**Authors:** Murphy De Meglio, Eloah S. De Biasi, Peter Breunig, Michael Candlish, Christina Sauerland, Stefan Günther, Haruya Kawase, Blanca Peguera, Christine Bohnstaedt, Peter R. Nilson, Jiong Hu, Susanne Hille, Oliver Müller, Amparo Acker-Palmer, Bruce D. Hammock, T Michael Underhill, Stephan Junek, Stefan Offermanns, Ingrid Fleming, Jasmin K. Hefendehl

**Author notes:** **Corresponding author:** Prof. Dr. Jasmin Hefendehl, Max-von-Laue Str. 13, Frankfurt am Main, Germany.

## Abstract

Recent advances in anti-amyloid therapies for Alzheimer’s disease have been promising, but they have also highlighted critical challenges, including increased vascular complications, such as amyloid-related imaging abnormalities. Emerging evidence suggests that the soluble epoxide hydrolase may be a promising therapeutic target due to the involvement of sEH-derived diols in inflammation, oxidative stress, and vascular destabilization.

APPPS1 mice were crossed with an inducible soluble epoxide hydrolase knock-out mouse line. The knock-out was induced before onset of amyloid deposition, and then the mice were analyzed using histological, molecular, and RNA sequencing techniques.

Here, we identify astrocytic soluble epoxide hydrolase as a key mediator of vascular instability in Alzheimer’s disease. Targeted astrocyte-specific deletion of soluble epoxide hydrolase in APPPS1 mice dramatically mitigated vascular changes, reducing the vascular amyloid burden by 67.95% and preserving VE-cadherin architecture. Importantly, vasomotion was markedly impaired in the Alzheimer’s disease model and was preserved in soluble epoxide hydrolase-deficient animals. Transcriptomic profiling of vasculature in APPPS1xsEHΔ^AC^ mice revealed upregulated expression of genes critical for neurovascular protection.

These findings identify soluble epoxide hydrolase as a central regulator of neurovascular dysfunction and underscore its therapeutic potential in increasing vascular stability in Alzheimer’s disease.

## Introduction

As the population ages, the incidence of Alzheimer’s disease (AD) is increasing, yet because of a lack of knowledge about disease pathogenesis there are no early-intervention therapies available. There is, however, agreement that vascular defects and dysfunction of the blood-brain barrier (BBB) are early events in the onset of AD^1–5^. Despite extensive research into amyloid-β (Aβ) pathology and cerebral amyloid angiopathy (CAA), the molecular mechanisms underlying the breakdown of the BBB in AD remain elusive.

In the cerebral circulation, there is a close association between endothelial cells, pericytes and astrocytes which make up the neurovascular unit (NVU), and there is evidence to implicate pericyte loss with compromised BBB integrity in AD patients^6,7^. One pathway that has been linked to pericyte loss and the breakdown of the blood-retinal barrier in diabetic mice and humans is linked to the soluble epoxide hydrolase (sEH). This enzyme can generate fatty acid diols such as 19,20-DHDP that can essentially dissolve endothelial cells as well as endothelial cell-pericyte junctions^8^. Importantly, sEH expression is elevated in models of vascular cognitive impairement^9,10^, small vessel disease^11^, type 2 diabetes^12^, and AD^13,14^. Studies have shown that inhibiting sEH can reduce neuroinflammation, lower parenchymal amyloid burden, enhance functional hyperemia, and improve cognitive function in AD models^13–15^. All of these reports suggest that targeting sEH may be a promising approach to delay key pathological features of AD^11,13–16^. Given the strong association between sEH and vascular dysfunction, this study set out to determine, 1) whether sEH disrupts astrocyte-pericyte communication in AD in a manner similar to that observed in diabetic retinopathy, 2) how astrocyte-specific sEH inhibition reduces parenchymal amyloid, as well as vascular amyloid deposition, which could pave the way for transformative advancements in AD treatment, and 3) how genetic ablation of astrocytic sEH impacts vasomotion in AD, with a particular focus on CAA.

Currently, many patients are excluded from current anti-amyloid therapeutic interventions due to cerebrovascular risk factors, such as CAA, which increase the risk of amyloid-related imaging abnormalities (ARIA)^17^. In light of the current ongoing amyloid trials, these findings underscore astrocytic sEH as a central regulator of vascular degradation in AD and reveal its inhibition as a promising therapeutic avenue for preserving neurovascular unit integrity and mitigating Aβ-driven neuropathology.

## Materials and methods

### Mouse Lines

All experimental procedures and husbandry were carried out in accordance with regulations of the state of Hessen (Germany), and by the animal welfare committee of Johann Wolfgang Goethe-Universität Frankfurt am Main. Mice had unrestricted access to food and water. All Cre systems were induced by 0.02g/ mL(5 µl /g) of tamoxifen via intraperitoneal injection for five consecutive days.

### Genotyping

Genotypes were determined from ear or tail biopsies by alkaline lysis (50 mM NaOH, 95 °C, 45 min) followed by neutralization (1.5 M Tris-HCl, pH 8.8).

### Stereotaxic intracranial Injections of adeno-associated viral vectors

Stereotaxic injections were performed under fentanyl/midazolam/medetomidine anesthesia. Coordinates were defined relative to bregma (AP: −0.15 mm; ML: ±0.02 mm; DV: −0.15 mm). Virus was infused at 0.1 μl/min, and the needle was left in place for 10 min before withdrawal, as described previously^18^. Mice received postoperative analgesia (buprenorphine, 0.015 mg/ml for 3 days) and were monitored until sacrifice.

### Tissue preparation and immunohistochemistry

Mice were perfused with PBS followed by 4% paraformaldehyde (PFA). Brains were post-fixed for 2 hours, cryoprotected in 30% sucrose, and sectioned coronally. Free-floating sections were permeabilized, blocked, and incubated with primary antibodies overnight at 4 °C, followed by fluorophore-conjugated secondary antibodies. For VE-cadherin detection, tyramide signal amplification (SuperBoost™, Invitrogen) was used according to manufacturer instructions. Sections were mounted in Fluoromount-G.

### Cranial window surgery

To be able to visualize the brain vasculature in a living mouse, cranial windows were implanted on the hindlimb somatosensory cortex as described^19^.

Briefly, the mice were anesthetized with a combination of fentanyl/midazolam/medetomidine (10 µl/g), their heads were shaved, and the mice were positioned in a stereotaxic apparatus. Once the head was fixed in position the area was cleaned with povidone- iodine solution followed by Softasept N.

After the surgery, the animals were removed from the stereotaxic apparatus, injected with buprenorphine (6.66 μl/g) and antidote (10 μl/g). Mice were then allowed to recover in the cage on a warm surface.

### Generation of adeno-associated viruses

An adeno-associated virus serotype 9 (AAV9)-based delivery system was employed to overexpress sEH in vivo. The open reading frame (ORF) of human *EPHX2* (sEH; transcript variant 1) was PCR-amplified from a HepG2 cDNA pool and cloned into a single-stranded AAV (ssAAV) genome plasmid under the control of the astrocyte-specific minimal human glial fibrillary acidic protein promoter (gfaABC1D; Lee et al., 2008). To enable identification and tracking of transduced cells, an eGFP reporter was co-expressed from the same transcript in a bicistronic design, linking sEH and eGFP via a self-cleaving porcine teschovirus-1 2A (P2A) peptide. A construct expressing eGFP under the gfaABC1D promoter served as control.

For AAV vector production, HEK293T cells at low passage were transfected with the respective AAV genome plasmid and the adenoviral helper construct pDP9rs, encoding the capsid and helper factors. After 72 h, cells and culture media were collected, and recombinant AAV particles were isolated using iodixanol gradient ultracentrifugation. Purified vectors were filtered, concentrated, and their genomic titer determined by quantitative PCR as described previously^20^.

### Western blotting of whole brain homogenates

Brains were perfused with PBS and then homogenized in RIPA buffer. Proteins were separated by SDS-PAGE (Novex 10-20% Tricine Mini Protein Gels, Invitrogen, cat. no. EC66252BOX, Novex Tricine SDS Running Buffer, Invitrogen, cat. no. LC1675) and transferred onto nitrocellulose membranes with 0.2 µm pores (Whatman, Dassel, Germany), using the Mini Trans-Blot® Cell (Bio-Rad). Membranes were probed over night with specific primary antibodies as indicated and then with peroxidase-conjugated anti-rabbit or anti-mouse secondary antibodies for 1 hour at room temperature. Bound antibodies were visualized by enhanced chemiluminescence reagent (Millipore) and a ChemiDoc MP Imaging System (Bio-Rad). Quantification of band intensities were analyzed by the Image Lab software (Bio-Rad, version 6.0.1). Equal loading was assessed using antibodies to GAPDH.

### Image acquisition and data analysis

Immunohistochemistry samples were analyzed with FIJI software and Imaris software (Imaris software (v 9.7), Oxford Instruments). *In vivo* data were analyzed with FIJI and, posteriorly, with Jupyter Notebook (run Phyton script) or Matlab (version R2022a).

### 2-Photon microscope

A custom made 2-Photon microscope built according to the steps described elsewhere ^21^ was used for the *in vivo* experiments. A Chameleon Ultra II laser (Coherent) was used as well as an MP285A (Sutter Instrument Company) for motor control and ScanImage software (Vidrio Technologies) for image acquisition. Laser emission was detected by two photomultiplier tubes (Hamamatsu Photonics), a 695dcxxr primary and a T560LPXR secondary dichroic mirror, an ET525/50m-2P and an ET605/70m-2P bandpass filter (Chroma Technology Corporation). A head fixation system that allowed imaging of the same regions of interest based on consistent coordinates was used, as described^22^. Images were acquired at a laser power < 50mW.

### RNA spatial sequencing

Brains were embedded in OCT and frozen in isopentane. Sections (10 μm) were processed using the 10x Genomics Visium Spatial Gene Expression platform according to the manufacturer’s protocols. RNA integrity was verified (RIN ≥7) prior to library preparation. Libraries were sequenced and analyzed following standard workflows.

### sEH/GFAP Co-localization analysis

Cortical regions of 9-month-old APPPS1 mice were imaged using a confocal laser scanning microscope. The images were analyzed in Fiji. The images were despeckled and had the background subtracted by using the rolling ball method with 50 pixels. The plugin colocalization colormap (version 12_11_2019) was used to get the index of correlation

### sEH proximity to plaques

Laser scanning confocal microscopy images were used for analysis of sEH immunoreactivity in proximity to plaques. The images were analyzed using Imaris. A consistent threshold was set, and the distance of the sEH surface from the Methoxy-X04 surface was recorded and then binned according to instances in which the distances were observed.

### sEH proximity to CAA

Laser scanning confocal microscopy images were used for analysis of sEH immunoreactivity in proximity to CAA by using hFTAA to visualize the CAA on vessels. In Fiji, the polygon tool was used to draw a ROI around CAA deposits. The selection area was enlarged by 25-microns. Selection of both ROIs, the XOR feature was used to isolate the 25-micron region around the CAA. The intensity of sEH immunoreactivity was measured in this ROI. For non-CAA burdened controls, similar sized vessels were used to measure the amount of sEH immunoreactivity in 25-micron regions.

### Intravenous Alexa Fluor 555 Cadaverine injection

Mice were put into surgical plane of anesthesia by injection of ketamine and xylazine at 180 mg/kg and 10 mg/kg of body weight, respectively. 70 μL of Alexa Fluor 555 cadaverine (1 mg/ml) was injected into the retro-orbital venous sinus and circulated for 30 minutes before mouse was sacrificed by overdose of ketamine/xylazine. Mice were then perfused with 4% PFA. Please see Tissue Preparation.

### Plaque analysis

Dense-core and diffuse amyloid were segmented using Methoxy-X04 and hFTAA, respectively, with Fiji particle analysis.

### APP and Cadaverine analysis

Epifluorescence images of cortical regions were processed using Fiji imaging software. The polygon tool was used to select the tissue. The intensity of the channel was measured and the mean intensity was recorded. The mean of each mouse was recorded for analysis.

### Process length analyses

Epifluorescence or max projection of laser scanning confocal images were analyzed using Fiji software. A skeleton of the respective signal and the normalization signal were created using a skeletonizing script provided by Dr. Michael Candlish and Dr. Eloah De Biasi. An algorithm from “Robust Quantification of In Vitro Angiogenesis Through Image Analysis"^23^ was used in which the lengths are calculated from the middle of one pixel to the middle of another.

### Cortex covered area by vascular-associated components in aging and disease

Images of the cortex area were taken at the epifluorescence microscope. The thresholded area was then skeletonized^23^. The results provided the length of the vasculature and *Hic1/* tdTomato^+^ cell coverage post skeletonization and was further normalized by tissue area to determine vessel coverage in the tissue as well as overall *Hic1/* tdTomato^+^ coverage. A total of two images per mouse was analyzed when possible and the acquired values were averaged and used for statistics.

### Hic1/ tdTomato^+^ somata counting and process length analyses

The images used for the previous analyses were also processed for *Hic1/* tdTomato^+^ somata counting and process length quantification. For somata counting, the images were processed using FIJI, where Gaussian blur filter sigma 2, followed by background subtraction using the rolling ball method (50 pixels) was applied. Afterwards, somatas were manually counted with the cell counter tool.

### Vasomotion analysis

To analyze variations in vasomotion in CAA burdened vessels and WT healthy vessels, acquired images were opened in FIJI. As described elsewhere^24^, a line was drawn across the arteriole of interest in FIJI and saved as a ROI for further application. A collection of scripts kindly shared to our group by Dr. Leon P. Munting were then used to acquire the power spectral density of each of the analyzed vessels as well as the maximum amplitude of vasomotion, which was showed directly at Matlab and posteriorly copied into Prism for statistics.

### Vasomotion measurements

To assess the presence of vasomotion in CAA+ and healthy vessels, heterozygous and wildtype APPPS1 mice aged 9-16 months were used. Cranial windows were implanted as previously described. For region of interest (ROI) determination, animals were anesthetized with fentanyl/midazolam/medetomidine (10 µl/g) and tail vein injected with Alexa 633 Hydrazide and 1% fluorescein in sterile saline to visualize the brain vasculature. A Plan-Apochromat W 40x/1.0 DIC VIS/IR (Zeiss) objective was used for ROI imaging. Alexa hydrazide positive vessels were considered arterioles. For vasomotion detection, animals were anesthetized with a combination of Ketamine (1 μl/g) and Xylazine (0.5 μl/g), to avoid Isoflurane interferences on cerebrovasculature tone^24^. Animals were subsequently i.p injected with 1% fluorescein in sterile saline. ScanImage software was used for vasomotion detection where 256x256 images were acquired. A total of 1266 frames were acquired at a 4.22 Hz rate for a total period of 5 minutes ^24^. Vessels were imaged at 800 nm with a zoom of 2, and the red channel was excluded from the data attainment.

### Microbleedings

For this experiment, *Hic1^CreERT^*^2^*;Rosa26^LSL-tdTomato^* APPPS1 mice ranging from 12 to 23 months of age were used. The vasculature was visualized with 1% fluorescein and Aβ with Methoxy-X04, injected 24h prior to imaging. Capillaries chosen to be ablated were approximately 100 µm below the brain surface. Z-stack images (512x512, 1 µm step size, 2x zoom) of pre and post ablation capillaries were acquired with a Plan-Apochromat W 40x/1.0 DIC VIS/IR (Zeiss) at 800 nm (optimal excitation of the fluorescein signal), 900 nm and 1047 nm (optimal excitation of the tdTomato signal).

### Statistical analysis

All statistical analysis was done using GraphPad Prism 10 software. All specific tests used are state in the figure legends or in the main text. Outliers were excluded using the ROUT method using standard settings in GraphPad Prism 10 software. All data in the graphs are shown as mean of each animal ± SEM, unless otherwise stated.

## Results

### Vascular stability is impaired in APPPS1 mice

BBB dysfunction is an early pathological event observed in individuals with AD^1,2^, therefore, we assessed BBB integrity in APPPS1 mice following intravenous injection of Cadaverine. This revealed detectable and generalized vascular leakage in APPPS1 mice (Extended Fig. 1A,B). To further characterize the underlying mechanisms of BBB disruption, we analyzed the organization of endothelial junctional proteins within the capillary bed. This revealed a pathology-dependent disorganization of adherens junctions, marked by fragmented and discontinuous VE-cadherin staining along the vascular endothelium. Such alterations in junctional architecture are consistent with compromised barrier integrity and thus contribute to the observed Cadaverine extravasation (Extended Fig. 1C,D). To better visualize how the cells of the NVU are affected in AD, we crossed APPPS1 mice with a stromal progenitor cell (SPC) reporter line, *Hic1*-tdTomato^+^, as SPCs play an essential role in the repair of injured vessels^25^. To investigate vascular repair dynamics, targeted *in vivo* capillary ablation was performed by laser lesioning, and recovery was followed over 28 days.

This revealed that the recovery of vessel flow was markedly impaired in *Hic1*-tdTomato^+^ APPPS1 mice (Fig. 1A,B). This indicates that AD pathology hampers efficient vessel repair and recovery, consistent with our initial hypothesis that the BBB dysfunction negatively affects vessel recovery after acute injury.

**Figure 1.**
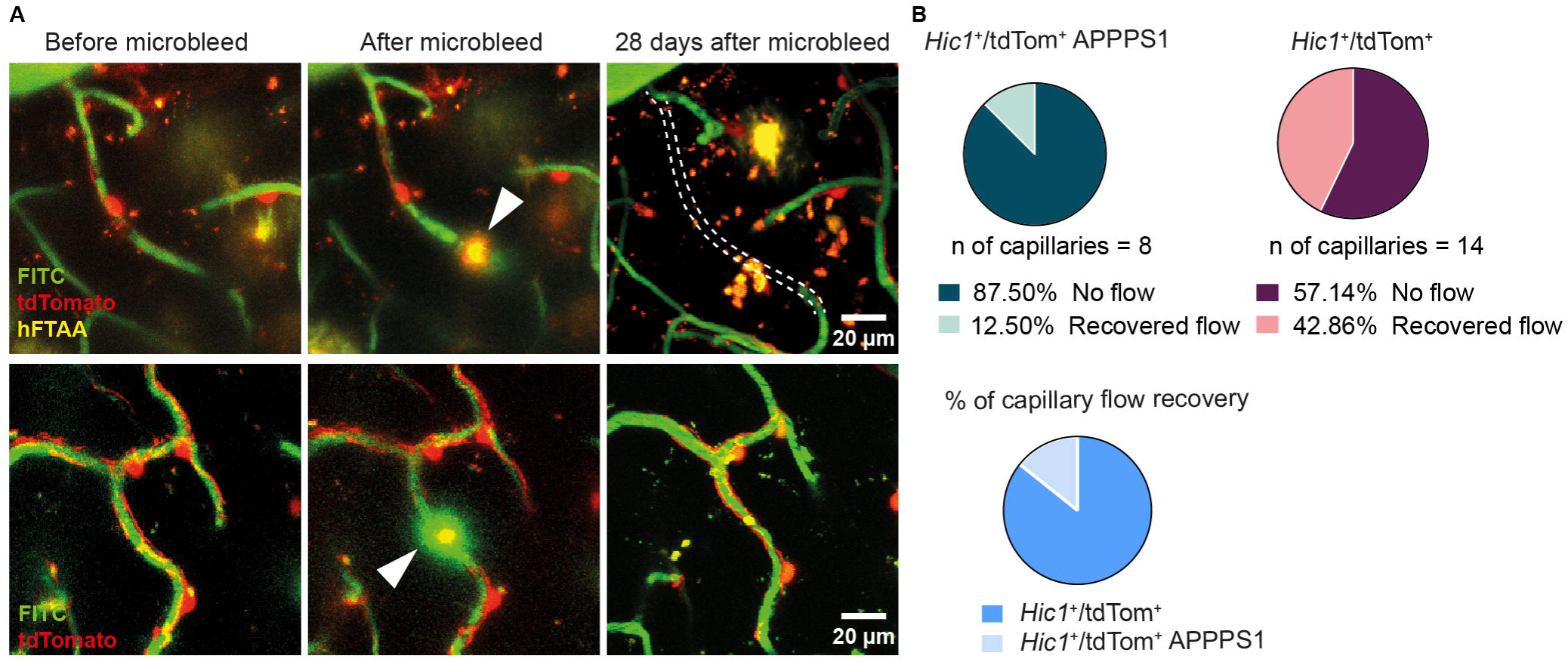
**Cerebral amyloidosis impairs capillary repair upon microbleeds**. (A) Representative pictures of a lost (upper image) and recovered (lower image) capillary, portraying the ROIs before, after and 28 days after the microbleed (indicated by arrowhead). Scale bar = 20 μm. (B) Pie charts displaying the percentages of lost and recovered vessels in *Hic1*-tdTomato^+^ APPPS1 and *Hic1*-tdTomato^+^ mice. *Hic1*-tdTomato^+^ APPPS1 animals have significantly less capillary recovery following microbleed, compared to *Hic1*-tdTomato^+^ mice.

Given the central role of *Hic1*-tdTomato^+^ SPCs as key regulators of BBB integrity, we hypothesized that alterations in this cell population might contribute to the vascular defects observed in AD. To address this, we observed *Hic1*-tdTomato^+^ cells in the APPPS1 mouse model. In the absence of AD, cortical areas of 8- and 20-month-old animals displayed stable *Hic1*-tdTomato^+^ cell coverage and somata numbers (Fig. 2A,B). In contrast, both parameters were significantly reduced in age-matched 8- and 20-month-old APPPS1 animals (Fig. 2C), indicating that AD pathology directly affects *Hic1*-tdTomato^+^ SPCs.

**Figure 2.**
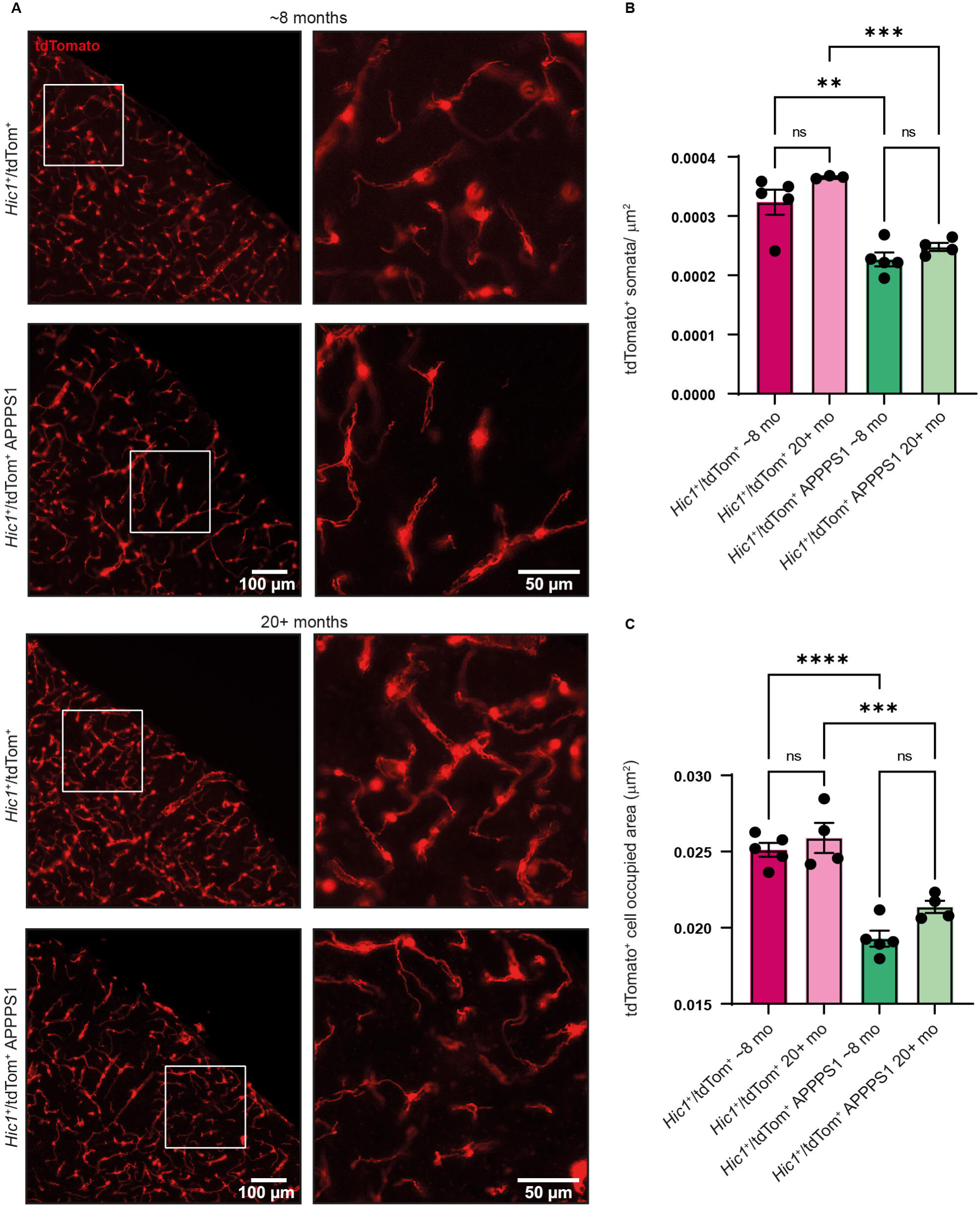
**SPC loss in the capillary bed in a model of cerebral amyloidosis. (**A) Representative images showing the *Hic1*-tdTomato^+^ cell signal in the mouse cortex in different ages, in control, and diseased mice. Zoomed in images highlight the reduction in cell coverage and *Hic1*-tdTomato^+^ somata. Scale bar = 100 μm; zoomed in scale bar = 50 μm. (B) Number of *Hic1*-tdTomato^+^ somata was unchanged in normal aging, showing a significant reduction in number in the disease model in both age groups. (C) Aging did not affect *Hic1*-tdTomato^+^ cell coverage, however, cerebral amyloidosis resulted in a significant reduction in SPCs in the cortical area (8 months *Hic1*-tdTomato^+^: n=5 mice, 8 months *Hic1*-tdTomato^+^ APPPS1: n=5 mice; 20 months *Hic1*-tdTomato^+^: n=4 mice; 20 months *Hic1*-tdTomato^+^ APPPS1: n=4 mice. Ordinary one-way ANOVA Šídák’s multiple comparisons; ** = p < 0.01, *** = p < 0.001 **** = p < 0.0001).

To assess whether the decrease in *Hic1*-tdTomato^+^ SPC somata and coverage was linked to vascular rarefaction, vascular density was calculated by analysing the length of vessels per area of brain. The results showed that vessel coverage of Podocalyxin expressing endothelial cells was unchanged in all animal groups (Extended Fig. 2A,B).

### Amyloid pathology drives robust sEH expression in cortical astrocytes

To further investigate the mechanisms underlying SPC loss, we examined the intercellular communication of astrocytes and SPCs, as both cell types play essential roles in maintaining vascular stability. Upregulation of astrocytic enzyme sEH has previously been implicated in vascular dysfunction and neurovascular unit instability^8,13^. Consistent with that, we observed elevated sEH expression in cortical areas from APPPS1 mice compared to wild-type controls, particularly in reactive astrocytes (Extended Fig. 3A-C). There was also a clear spatial association between cells with high sEH expression and Aβ deposits detected using Methoxy-X04 (Extended Fig. 3D,E). Given our focus on the vasculature, we examined the relationship between sEH expression and Aβ deposition along cerebral vessels. This revealed consistently stronger astrocyte sEH immunoreactivity close to vessels in APPPS1 mice in comparison to age-matched controls (Fig. 3A). The most striking finding was that sEH expression was particularly high around CAA-burdened (hFTAA) vessels (Podocalyxin) versus non-burdened vessels in the same animal (Fig. 3B,C), suggesting a direct link between CAA and sEH upregulation.

**Figure 3.**
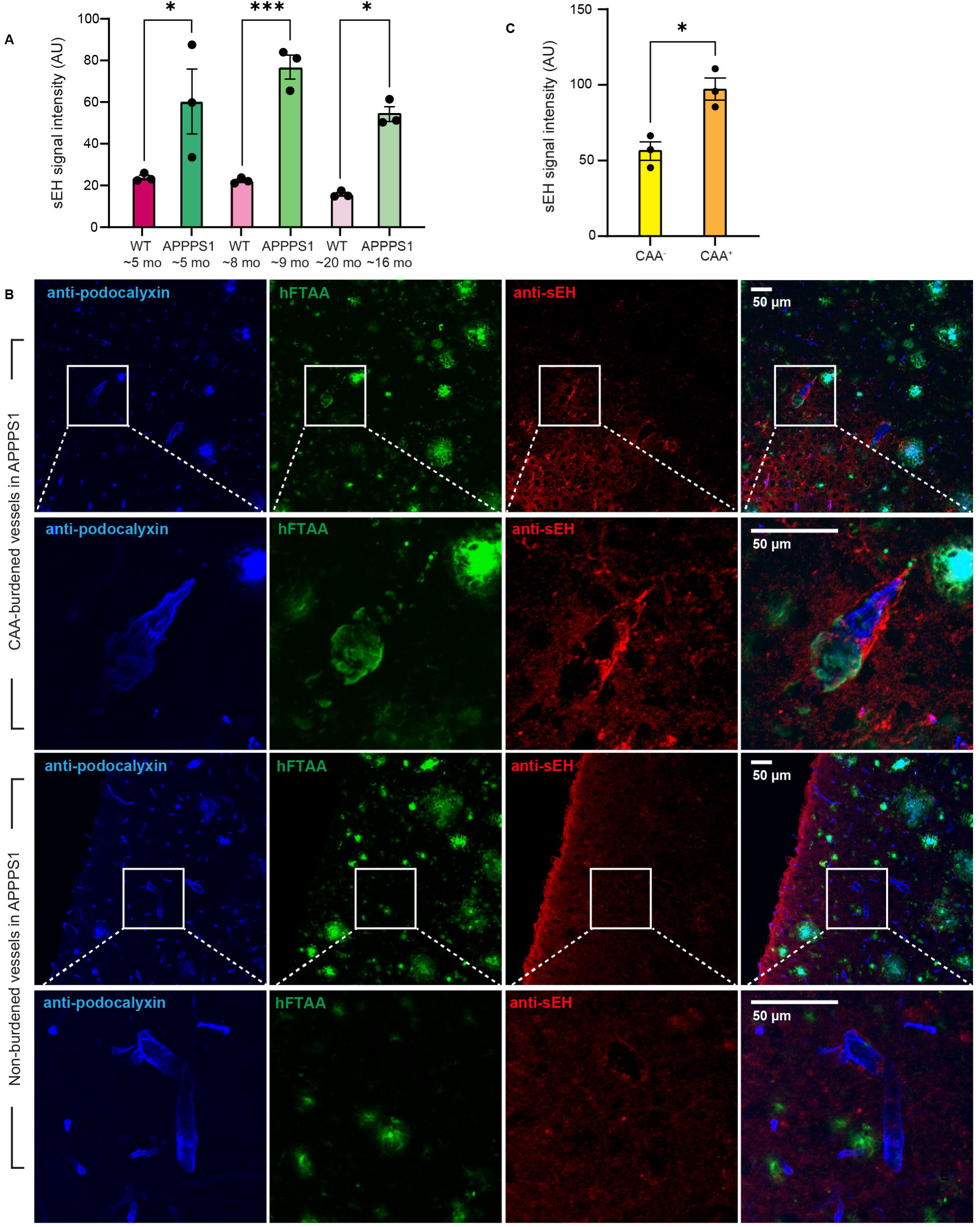
sEH expression increases around CAA-burdened vessels. (A) APPPS1 mice have higher sEH immunoreactivity around vessels compared to age-matched controls (n = 3, Ordinary one-way ANOVA; * = p < 0.05, *** = p < 0.001 (B,C) CAA-burdened vessels are associated with sEH immunoreactivity, when compared to CAA-negative vessels (n = 3, two-tailed unpaired t-test, * = p < 0.05).

### Vascular amyloidosis is associated with loss of arteriolar *Hic1*-tdTomato^+^ cells

Next, we sought to determine whether the observed *Hic1*-tdTomato^+^ SPC loss was spatially linked to CAA. When comparing *Hic1*-tdTomato^+^ SPC coverage on CAA burdened (CAA^+^) and unburdened arterioles (CAA^-^) we found that *Hic1*-tdTomato^+^ SPC numbers were significantly reduced on CAA^+^ arterioles, while *Hic1*-tdTomato^+^ SPC coverage was similar on CAA^-^ arterioles from APPPS1 mice compared to wild-type controls (Fig. 4A,B). To determine the physiological impact of arteriolar CAA accumulation, we assessed vasomotion in CAA^+^ and wild-type mice arterioles and detected a significant reduction in the maximum amplitude of vasomotion frequency in CAA^+^ arterioles compared to controls (Fig. 4C-E). Thus, vascular amyloidosis has a direct, negative effect on spontaneous vascular contractions. Together, our findings identified coincident Aβ-associated upregulation of sEH, loss of *Hic1*-tdTomato^+^ SPCs, and impaired vascular integrity.

**Figure 4.**
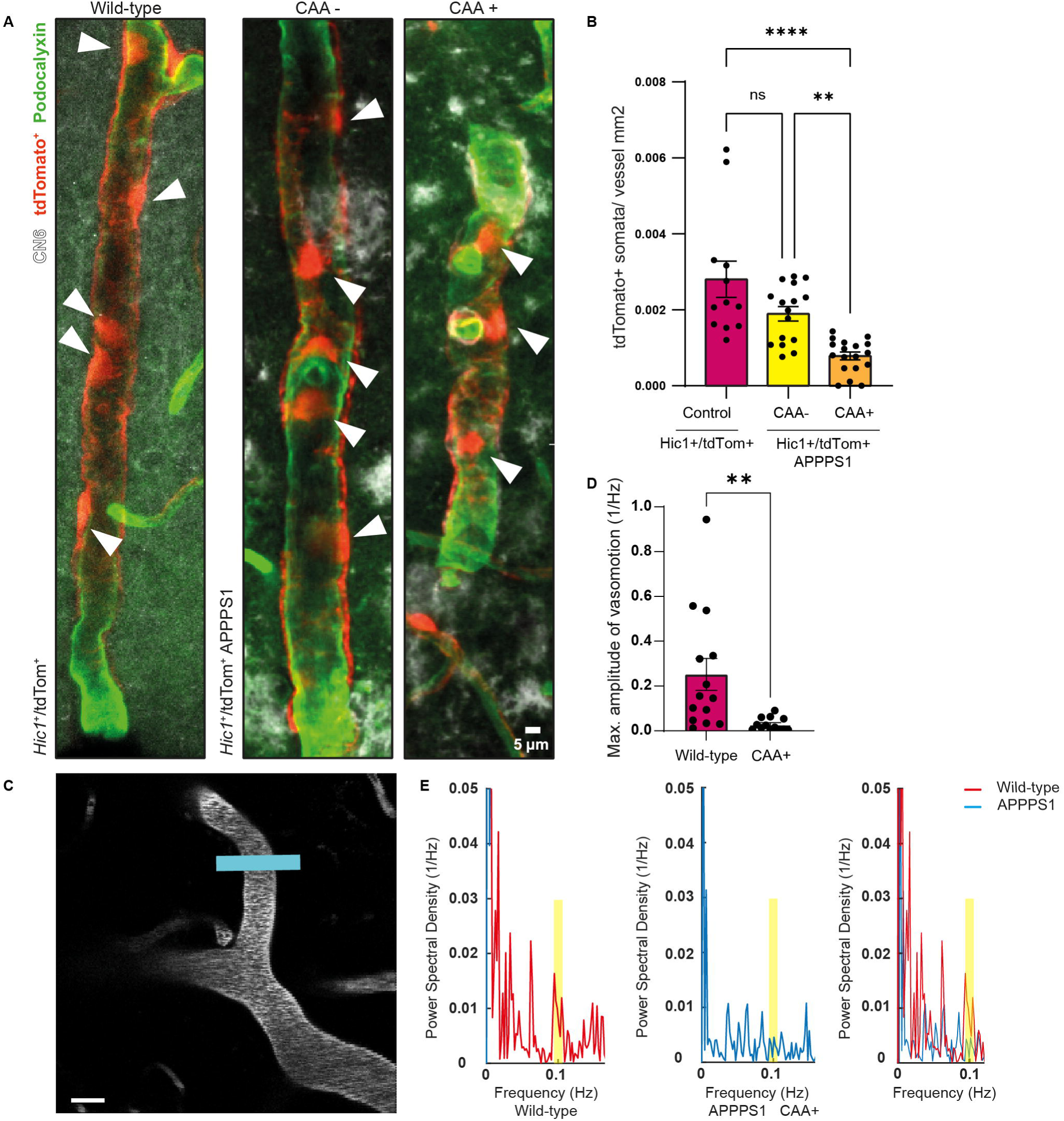
CAA associated with tdTomato+ cell loss at 8 months. (A) Representative images showing arterioles (green), *Hic1*-tdTomato^+^ cells (red) and CAA (white) in 8-month *Hic1*-tdTomato^+^ and *Hic1*-tdTomato^+^ APPPS1 CAA+ and CAA-. Scale bar = 5 μm. (B) CAA+ vessels have reduced *Hic1*-tdTomato^+^ somata coverage in comparison to CAA- vessels and control: *Hic1*-tdTomato^+^ (control): n = 12 vessels (5 mice), *Hic1*-tdTomato^+^ APPPS1 CAA-: n=16 vessels (5 mice), *Hic1*-tdTomato^+^ APPPS1 CAA+: n=18 vessels (5 mice); Ordinary one-way ANOVA Šídák’s multiple comparisons; ** = p < 0.01, **** = p < 0.0001. (C) Representative figure of one of the ROIs analyzed for vasomotion. Scale bar = 20 μm. (D) Significantly different maximum amplitude of vasomotion (0.1Hz) between wild-type and CAA+ vessels from APPPS1 animals (Wild-type: n = 14 vessels (4 mice), APPPS1 CAA+: n= 13 vessels (3 mice). Two-tailed unpaired t-test; ** = p < 0.01). (E) Representative power spectral density graphs from a wild-type and an APPPS1 mouse. Notice the smaller amplitude of the peaks around 0.1Hz in the APPPS1 mouse (yellow rectangle).

### Impact of astrocytic deletion of sEH in APPPS1 mice

To demonstrate cause and effect, and make a functional link between sEH and the vascular pathology characterized by SPCs, APPPS1 mice were crossed with inducible astrocyte-specific sEH knockout mice, generating APPPS1 x *Ephx2* flox x *Aldh1l1*-Cre/ERT2 (herein referred to as APPPS1xsEHΔ^AC^).

To determine whether astrocytic sEH contributes to the pathogenesis of AD, tamoxifen was administered to 4–5-week-old mice i.e., before the onset of Aβ deposition and amyloidosis was studied at 9 months of age. Given the distinct roles of dense core plaques and diffuse amyloid depositions in AD pathology^26^, we quantified both forms through histological analysis to assess potential differences in Aβ burden. Our analysis showed that astrocytic specific sEH deletion led to a dramatic (over 60%) reduction in dense core plaques as well as a more subtle, but still significant decrease in diffuse amyloid (Fig. 5A,B). Overall Aβ burden in the APPPS1xsEHΔ^AC^ mice was significantly reduced (Fig. 5C), indicating that the deletion of astrocytic sEH had a robust influence on Aβ deposition in this model. Most importantly, beyond parenchymal plaques, CAA burden on vessels was dramatically reduced by (over 65%) in APPPS1xsEHΔ^AC^ when compared to APPPS1 mice (Fig. 5D).

**Figure 5.**
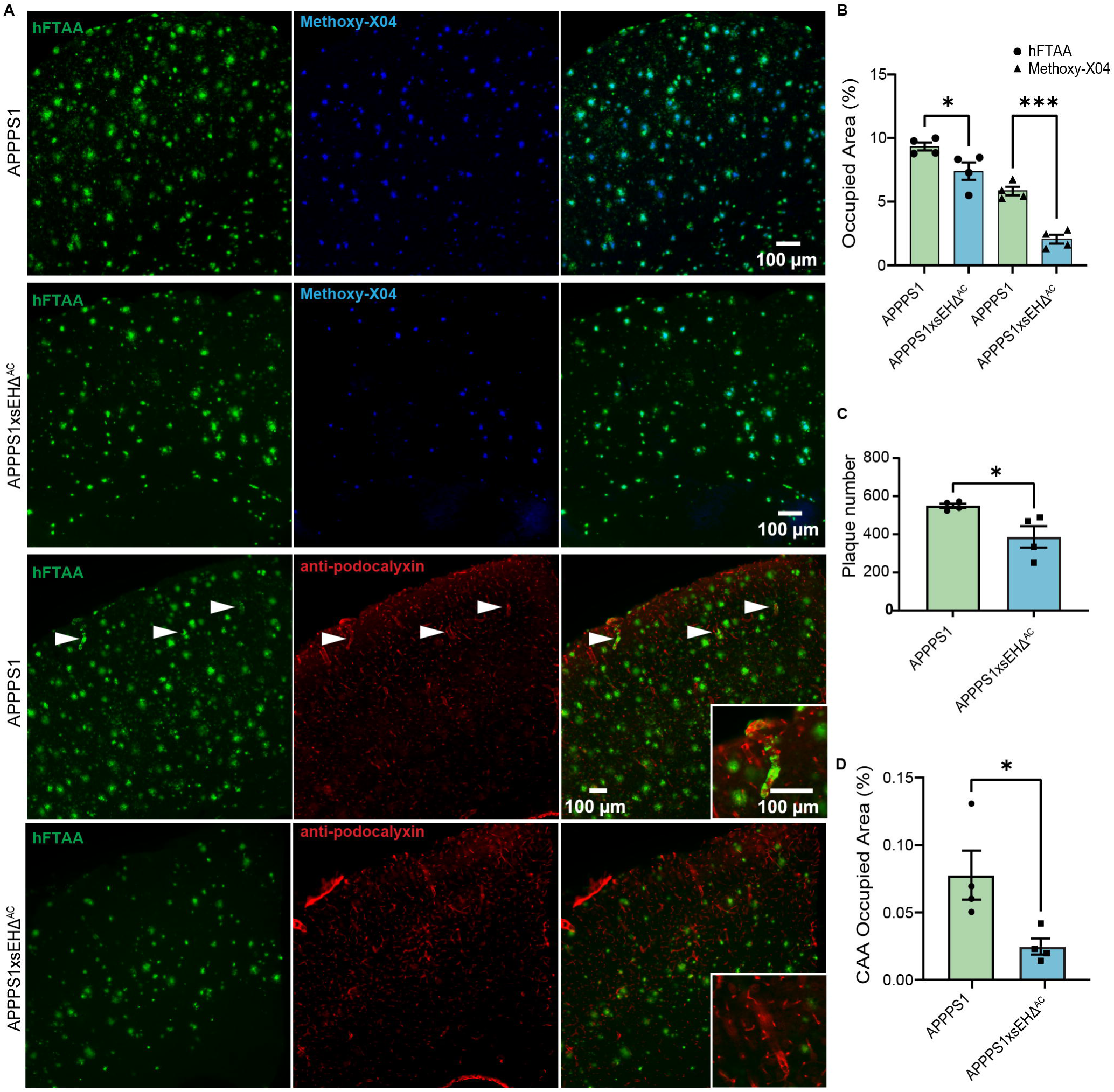
Knocking sEH out in APPPS1 mice has a neuroprotective effect by significantly reducing amyloid plaque burden and CAA. (A) Representative images of amyloid burden reduction between APPPS1xsEHΔ^AC^ and APPPS1 mice. (B) When crossbreeding an astrocyte-specific sEH knock out mouse with the APPPS1 mouse, a significant reduction in diffused amyloid (hFTAA, green) and dense amyloid (Methoxy-X04, blue) can be observed (n = 4, Ordinary one-way ANOVA, * p = < 0.05, **** p = < 0.0001). (C-D) Not only is there a significant reduction in plaque covered area, but there is also a significant reduction in overall plaque numbers (n = 4, two-tailed unpaired t-test, * = < 0.05). (E) Most significantly, there is a decrease in CAA occupied area (n = 4, two-tailed unpaired t-test, * p = < 0.05).

Aβ plaques are closely associated with dystrophic neurites, which are indicators of neuronal dysfunction and degeneration^27^. Given the observed reduction in Aβ burden, we next investigated whether sEH deletion could mitigate neuronal injury. Indeed, we detected a 30.11% reduction in amyloid precursor protein (APP), a marker for neuronal injury, around amyloid plaques in APPPS1xsEHΔ^AC^, indicating that astrocyte-specific deletion of sEH has profound therapeutic effects on both amyloid burden and amyloid-driven neurotoxicity (Extended Fig. 4A-C).

To further dissect the pathway by which astrocyte-specific sEH deletion can lead to a decrease in Aβ load, we analyzed APP processing, however, we found it to be unaffected (Extended Fig. 5A,B). We then looked at LRP1 because it is a main transporter responsible for the efflux of Aβ across the BBB, and its upregulation has been associated with enhanced Aβ clearance from the brain^28^. Given its involvement in Aβ metabolism and vascular health^29–31^, we hypothesized that the reduction in Aβ could be caused by increased levels of LRP1 expression. Using whole-brain lysates, we were able to detect a significant increase in LRP1 expression in APPPS1xsEHΔ^AC^ mice (Extended Fig. 5C), indicating that the observed reduction in Aβ burden is at least partially caused by increased clearance via LRP1.

### Deletion of astrocyte-specific sEH increases vascular integrity in APPPS1 mice

Our findings suggested that the Aβ-induced upregulation of the sEH in astrocytes and CAA leads to *Hic1*-tdTomato^+^ SPC loss and vascular dysfunction/BBB breakdown. To test this, we assessed relative VE-cadherin concentration^32^, a key protein in maintaining vascular integrity.

While APPPS1 mice showed a significant 32.95% reduction in relative VE-cadherin concentration, compared to wild-type controls, APPPS1xsEHΔ^AC^ mice exhibited only a 2.27% reduction (Fig. 6A,B). These results reveal a substantial preservation of vessel stability in APPPS1xsEHΔ^AC^ mice, suggesting that astrocytic sEH deletion provides a protective effect, despite the presence of Aβ pathology.

**Figure 6.**
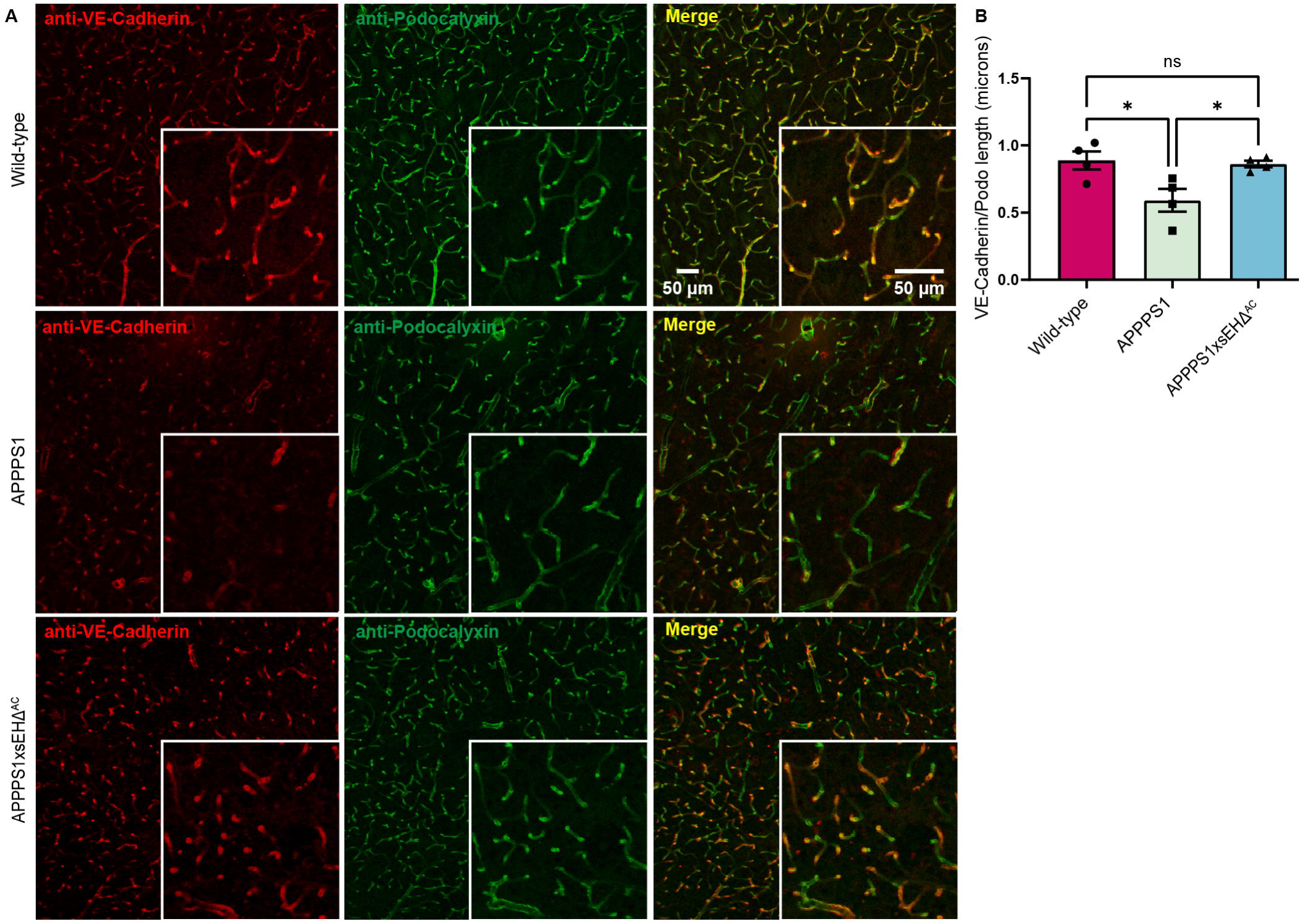
Knocking out sEH increases vascular integrity in AD mice. (A-B) Relative VE-cadherin concentration has a significant reduction in APPPS1 mice, however, there is a pronounced protective effect on vascular integrity when sEH is knocked out, as evidenced by the absence of discernible differences between the knockout model and the wild-type mouse (n = 4, Ordinary one-way ANOVA, * p = < 0.05).

### Viral overexpression of sEH induces BBB disruption and vascular cell loss

To provide a stronger mechanistic link between sEH and BBB dysfunction, and to exclude additional effects of AD pathology, a viral vector was used to locally overexpress the sEH in astrocytes of wild-type mice. The AAV9-ssgfaABC1D-sEH-2PA-EGFP viral vector used contained a gfaABC1D promotor region to facilitate the selective targeting of astrocytes. Controls animals underwent identical procedure and were treated with a control AAV9-ssgfaABC1D-EGFP virus. This control virus shared the same promoter and reporter gene, so that the target regions could be identified, and the assessment of mechanical disruption of the injection could be accounted for in the analysis.

The bicistronic expression of sEH and eGFP facilitated the visualization of sEH overexpression. In the viral target region, a 77.32% local reduction of relative VE-cadherin concentration was observed, when compared to the contralateral region (Fig. 7A,B). This magnitude of reduction was not detected in sham mice injected with control virus, where there was only a 5.44% reduction in relative VE-cadherin concentration between the target region and the contralateral side (Extended Fig. 6A,C). Collectively, the preservation of VE-cadherin in the APPPS1xsEHΔ^AC^ mice and the localized reduction of VE-cadherin in wild-type mice transduced to overexpress sEH in localized areas provide compelling evidence that sEH significantly contributes to compromised BBB integrity in AD.

**Figure 7.**
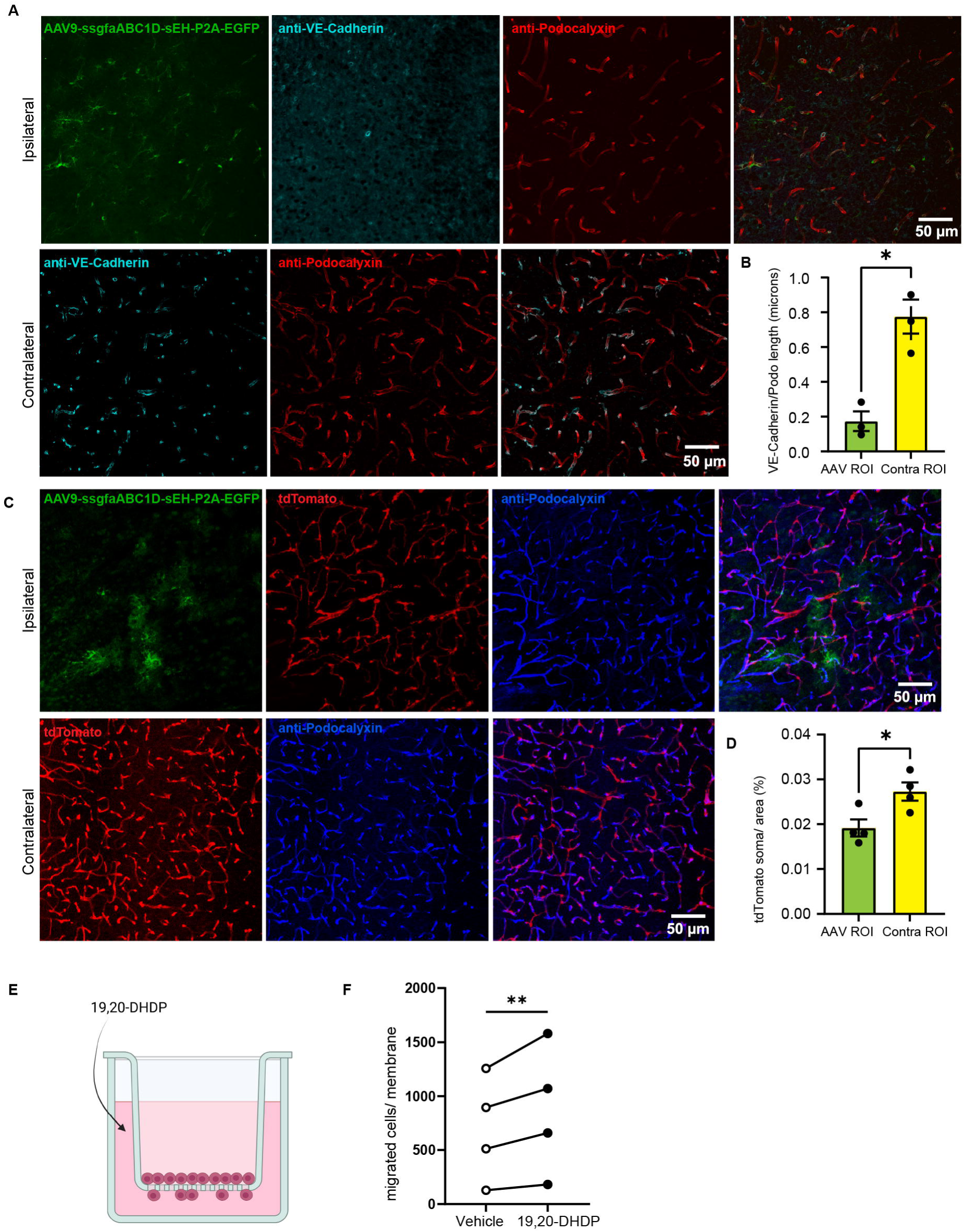
Overexpression of sEH is associated with BBB disruption. (A-B) Relative VE-cadherin (cyan) concentration was reduced within the viral region (VR) (green) (n = 3, one-tailed paired t-test, * p = < 0.05). (C-D) There is *Hic1*-tdTomato^+^ cell loss (red) on vessels (blue) in VR (n = 4, one-tailed paired t-test, * p = < 0.05). (E) Schematic of migration assay used with *Hic1*-tdTomato^+^ cells. (F) Isolated *Hic1*-tdTomato^+^ cells were treated with 3 μM of 19,20-DHDP for 16 hours, causing a significant amount of migration *in vitro* compared to vehicle control (n = 4, two-tailed ratio paired test, ** p = < 0.005).

In line with our observations of SPC loss in APPPS1 mice, we also detected a significant loss in *Hic1*-tdTomato^+^ cells within the viral target region in wild-type mice, compared to the contralateral side (Fig. 7C,D). In regions close to sEH expressing cells, there was a 29.63% reduction of *Hic1*-tdTomato^+^ compared to the contralateral side, but only a 4.59% reduction of *Hic1*-tdTomato^+^ in the viral target region in mice injected with control virus (Extended Fig. 6B,D). This result underscores a striking correlation between sEH upregulation and vascular dysfunction.

To determine a causal relationship between sEH modulation and the resultant loss of *Hic1*-tdTomato^+^ cells in both the APPPS1 model and the viral overexpression experiments, we conducted a migration assay using 19,20-DHDP, a diol produced generated by the sEH^33,34^ that can stimulate pericyte migration^8^. Previously published data indicate that 19,20-DHDP influences pericytes and perivascular fibroblasts, which make up 60.8% and 17.8% of our *Hic1*-tdTomato^+^

SPC population, respectively, by inducing a morphological shift towards a migratory phenotype ^8,25^. Indeed, treating *Hic1*-tdTomato^+^ SPCs with 19,20-DHDP for 16 hours (Fig. 7E,F) had a significant impact on the migration of SPCs *in vitro.* These results allowed us to conclude that sEH activity can precipitate *Hic1*-tdTomato^+^ SPC loss, in the absence of any underlying disease state or existing pathology/pre-pathology.

### Transcriptomic profile of APPPS1xsEHΔ^AC^ vessels reveal vascular resilience

Next, spatial RNA sequencing was performed to gain deeper insight into the transcriptomic differences in APPPS1xsEHΔ^AC^ close to the vasculature. Our findings reveal a coordinated upregulation of genes associated with vascular stability and neuronal protection. Of the differentially expressed genes, we found four that were particularly important for vascular stability, neuroprotection, and immune response—Glial cell line-derived neurotrophic factor (*Gdnf*)^35^, Sushi, Von Willebrand Factor Type A, EGF And Pentraxin Domain Containing 1 (*Svep1*)^36,37^, Protocadherin 12, or VE-cadherin-2 (*Pcdh12*)^38^, and chemokine ligand 1 (*Cxcl1*)^39^. All of these genes were upregulated in the vessels from APPPS1xsEHΔ^AC^ versus APPPS1 mice (Fig. 8A,B). Specifically, *Gdnf;* which encodes GNDF, one of the most potent neuroprotectants for dopaminergic neurons^40^, was upregulated in APPPS1xsEHΔ^AC^ vessels (Fig. 8B). This finding was particularly relevant as GDNF also contributes to the maintenance of endothelial barrier function^41,42^, suggesting a dual role in both neuronal and vascular protection. The upregulation of *Pcdh12,* which encodes for VE-cadherin-2, further contributes to vascular integrity through its superior aggregation properties compared to conventional VE-cadherin^43^. *Svep1* is involved in immune response regulation by modulating vascular permeability through endothelial remodeling processes^44^. Additionally, *Cxcl1* promotes endothelial cell survival and supports barrier function by stimulating endothelial repair mechanisms through enhanced endothelial cell-cell interactions and cytoskeletal reorganization^45^. Beyond its vascular effects, *Cxcl1* coordinates inflammatory response via recruitment of neutrophils and other immune cells^46^. The upregulation of these genes suggests that the deletion of sEH in astrocytes may trigger an adaptive or protective response in the vasculature, restoring vascular integrity through immune modulation.

**Figure 8.**
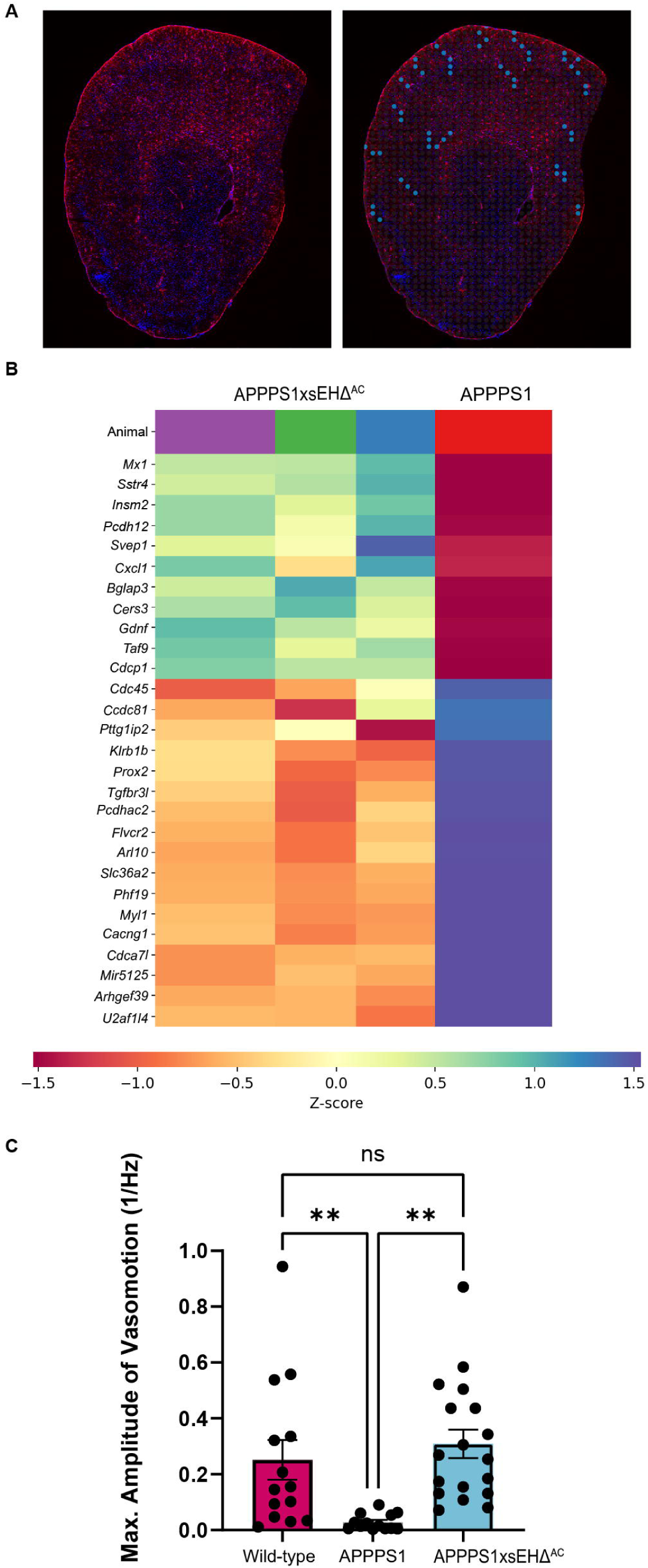
Transcriptomic profile of vessels altered in APPPS1xsEHΔ^AC^ mice compared to APPPS1 mice. (A) Representative image of vessel selection using RNA spatial sequencing. (B) Heat map of spatial sequencing data shows that DEGs between vessels of APPPS1 mice and vessels of APPPS1xsEHΔ^AC^ mice. (C) Vasomotion is preserved in APPPS1xsEHΔ^AC^ mice when compared to APPPS1 and wild-type mice. Wild-type: n = 14 vessels (4 mice), APPPS1: n= 18 vessels (5 mice), APPPS1xsEHΔ^AC^: n= 18 vessels (6 mice) Two-way ANOVA Tuckey’s multiple comparisons; ** = p < 0.01).

### Rescue of vasomotion in APPPS1xsEHΔ^AC^ mice

Remarkably, assessment of cerebrovascular dynamics in APPPS1xsEHΔ^AC^ mice revealed a striking preservation of vasomotion (Fig. 8C). While control APPPS1 mice exhibited profound impairment in vasomotion, the genetic ablation of astrocytic sEH appeared to fundamentally improve this process. Even though we observed CAA occurrence on some vessels at later ages, APPPS1xsEHΔ^AC^ mice maintain superior functional integrity and resilience compared to APPPS1 mice. These findings highlight the potential of sEH inhibition as a therapeutic strategy capable of persevering neurovascular coupling even in the context of established amyloid pathology.

## Discussion

Our findings reveal that BBB dysfunction in AD is closely linked to sEH-driven *Hic1^+^*cell loss and impaired vasomotion, particularly in vascular regions harboring Aβ deposits. Accordingly, we observed that CAA—characterized by the accumulation of Aβ in vessel walls —strongly correlates with these pathological changes, underscoring that localized vascular Aβ burden directly contributes to BBB impairment in AD. Furthermore, we reveal a spatially regulated mechanism through which Aβ associated sEH upregulation around CAA drives BBB breakdown through loss of *Hic1^+^*cells and VE-cadherin reduction, causing pathological structural alterations to the vasculature.

In APPPS1 mice, we observe chronic vascular issues, such as vascular leakage, unstable BBB, and disease-related reduced *Hic1^+^*cell coverage similar to those seen in patients with AD^47^. Moreover, we observed robust sEH expression in the cortical area that is not observed in the wild-type condition. Notably, sEH upregulation is most pronounced around CAA-burdened vessels, where it coincides with a marked reduction of *Hic1^+^*cells. Given that *Hic1^+^*cells serve a primary role in maintaining BBB integrity, this spatial relationship suggested a potential casual mechanism between sEH activity and vascular dysfunction.

We speculate that lipid metabolism mediated by sEH may impact vascular stability, as shown previously in models of diabetic retinopathy^8^. Specifically, the metabolic byproduct 19,20-dihydroxydocosapentaenoic acid (19.20-DHDP) has been shown to displace VE-cadherin from endothelial cell junctions via incorporation into lipid rafts, ultimately leading to pericyte loss. Consistent with this hypothesis, our *in vitro* experiments revealed that 19,20-DHDP induces migration of *Hic1*^+^, offering mechanistic insight into our reported findings. AD affects multiple cell types, making it challenging to isolate a single mechanism of action. To circumvent the potential confounding effects of AD pathology on our target cells *in vivo*, we employed viral overexpression of astrocytic sEH in wild-type mice to investigate its specific role in the absence of disease-related influences. With this experiment, we demonstrate that local sEH overexpression alone was enough to lead to a localized loss of *Hic1*^+^ cells and disruption of the junctional protein VE-cadherin, contributing to BBB dysfunction. Hence, these findings directly implicated sEH activity in vascular destabilization independent of AD pathology. As our data demonstrate, increased sEH expression during AD progression is spatially associated with amyloid deposition, whether in the parenchyma or along vessel walls, thereby initiating the described mechanistic pathway. We thus hypothesized that suppressing sEH activity has the potential to rescue the observed vascular dysfunction in APPPS1 mice. To test this, we generated an astrocyte specific sEH knock out model crossed with APPPS1 mice (APPPS1xsEHΔ^AC^). This data indeed showed that genetic ablation of astrocytic sEH alone is sufficient to preserve vascular stability, even in the presence of ongoing AD pathology. This stabilization allows for the sustained integrity of the BBB, evident by the preservation of VE-cadherin. Our findings confirm that the observed pathological alterations are driven by sEH activity and not by other potential AD-related mechanisms or downstream effects, and that targeted removal of sEH can effectively rescue vascular function despite AD pathology.

These findings provide compelling evidence that astrocytic sEH is a key regulator of vascular integrity, underscoring the promise of it being a therapeutic target for mitigating vascular dysfunction. Even more compelling is the association of reduced Aβ burden in APPPS1xsEHΔ^AC^ mice, that we show is due to enhanced clearance via transport across the BBB by LRP1, which has increased protein levels in APPPS1xsEHΔ^AC^ mice^48^. However, in addition to the increase in potential transport of Aβ across the BBB via LRP1, we cannot rule out the contribution of additional mechanisms influencing Aβ dynamics, such as altered glial activity, or changes to other perivascular clearance pathways.

In addition to the vascular alterations, we show that sEH is involved in inflammation-related pathological processes given its co-localization with reactive astrocytes. sEH metabolizes EETs, key lipid mediators which stunt pro-inflammatory pathways and cytokine-induced endothelial activation^33^. Beyond depleting these protective molecules, sEH upregulation has been associated with increased expression of adhesion molecules such as VCAM-1 and ICAM-1, which enhances leukocyte adhesion and infiltration of immune cells.^33,49^ While this response is beneficial in a healthy condition - facilitating immune surveillance and tissue repair - in diseases like AD, where we show that injury response is blunted, this can impede normal cellular functions and contribute to the disease progression.

Reduced relative VE-cadherin concentration and *Hic1*^+^ cell loss compromises BBB integrity and could contribute to chronic neuroinflammation and, in turn, Aβ accumulation by further impairing its clearance from the brain and allowing peripheral inflammatory factors to enter, actively driving deposition. Our results show that sEH inhibition is a viable therapeutic strategy, not only due to the reduction of Aβ burden shown in this study, but also because it promotes the stability of the neurovascular unit. Most interestingly, vasomotion analyses revealed that genetic ablation of astrocyte-specific sEH maintained proper vascular function when compared to APPPS1 mice, demonstrating that sEH deletion allows for functional protection to the cerebral vasculature despite the AD pathology. Notably, although some CAA coverage is observed in older APPPS1xsEHΔ^AC^ mice, this does not appear functionally relevant as vasomotion is restored. This preservation of vascular function likely stems from increased LRP1 expression, which enhances amyloid beta clearance. These findings may reflect a temporal shift of CAA onset in APPPS1xsEHΔ^AC^ mice, or possibly protective alterations in Aβ maturation state that preserve vascular activity. We and others have found that CAA predominantly compromises arterioles^50^, the vessels responsible for vasomotion (Extended Fig.7A,B). Notably, we observe a clear correlation between vasomotion dysfunction and CAA occurrence. Given that vasomotion modulates cortical blood flow at resting state by about 20%, these findings carry exceptional importance for the impact on brain health^51^.

The therapeutic implications of these findings extend beyond basic neurovascular mechanisms and offer potential solutions to critical limitations in current AD treatment paradigms. Currently, anti-amyloid antibody therapies present a promising future in AD management, however, their application is substantially restricted by safety concerns, especially among patients with cerebrovascular risk factors, such as the presence of CAA or APOE Ɛ4 alleles due to their increase risk of Amyloid-Related Imaging Abnormalities (ARIA), which can cause headaches, confusion, seizures, and even hemorrhages^52–56^. Our results show that sEH inhibition preserves vascular integrity and function even in vessels affected by CAA. By pharmacologically targeting sEH, it may be possible to fortify the NVU against the destabilizing effects of anti-amyloid antibodies. Such an approach could potentially expand the therapeutic window for treatment interventions to include previously excluded populations. Our study provides strong evidence that administering an sEH inhibitor in conjunction with current Aβ amyloid interventions could aid in the efficiency of the drugs, while also reducing the risk of ARIA through increasing the integrity of NVU.

In summary, we have identified a pivotal role of sEH in the AD pathology. We show that genetic ablation of astrocyte-specific sEH in our AD model preserves vascular stability, mitigates Aβ burden, and alleviates associated neuronal injury. Conversely, we have found that, in otherwise healthy mice, sEH overexpression causes a localized reduction of VE-cadherin, as well as *Hic1^+^* cell loss. These findings underscore the importance of sEH in NVU dynamics, and therefore, the importance of regulating sEH activity in maintaining vascular homeostasis. This strongly highlights sEH inhibition as a potential early-intervention therapy for AD, and that it could also enhance the effectiveness of current therapies by improving BBB integrity and NVU resilience, potentially lowering the risk of ARIA.

## Data availability

Any additional requests for data should be addressed to Prof. Jasmin Hefendehl.

