## Supplemental Figures for "Soluble epoxide hydrolase in Alzheimer’s disease drives neurovascular dysfunction"

### Supplementary material

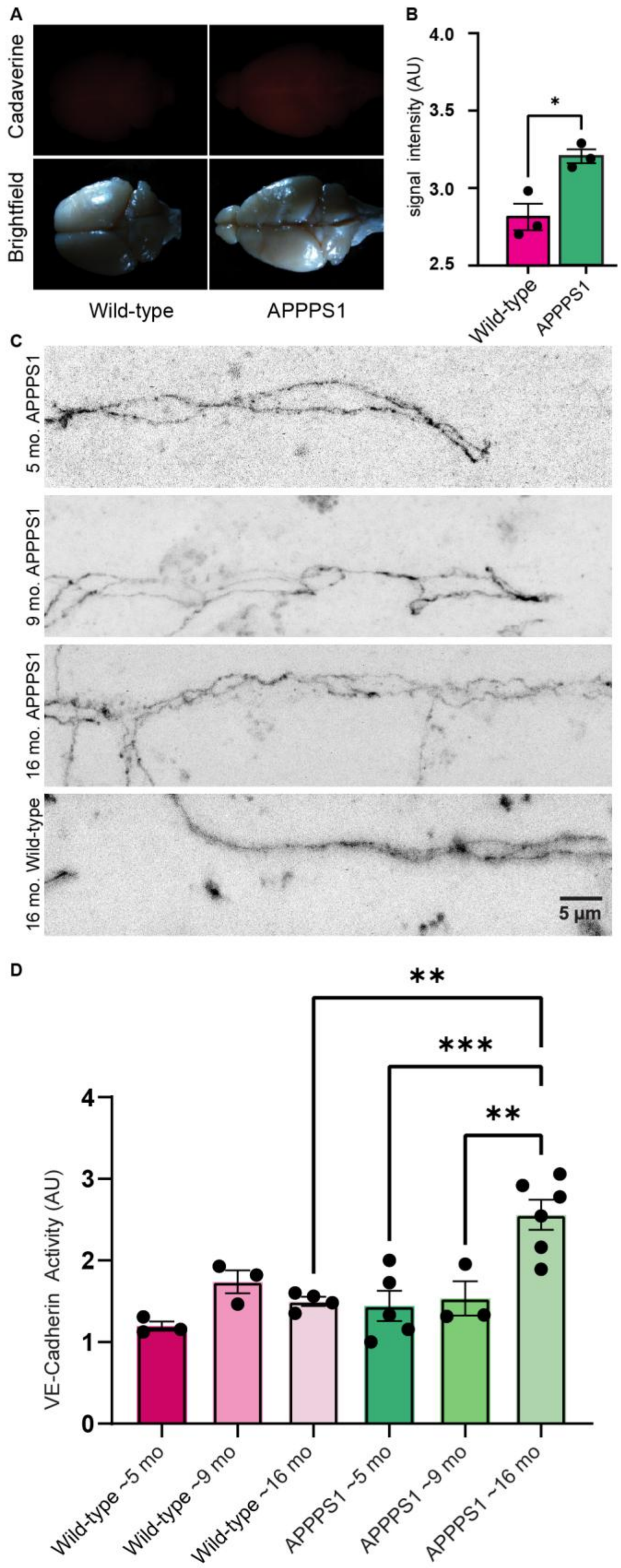
A
Cadaverine
Brightfield
Wild-type
APPPS1
B
4.0
3.5
3.0
2.5
signal intensity (AU)
*
Wild-type
APPPS1
C
5 mo. APPPS1
9 mo. APPPS1
16 mo. APPPS1
16 mo. Wild-type
5 µm
D
**
***
**
4
3
2
1
0
VE-Cadherin Activity (AU)
Wild-type ~5 mo
Wild-type ~9 mo
Wild-type ~16 mo
APPPS1 ~5 mo
APPPS1 ~9 mo
APPPS1 ~16 mo

**Extended Data Figure 1. Vascular stability is impaired in APPPS1 mice.** (A-B) More leakage was observed in APPPS1 mice compared to wild-type controls (n = 3, two-tailed unpaired t-test, * p = < 0.05). (C-D) Discontinuous VE-cadherin organization occurred in a pathogenic-dependent manner, when compared to age-matched controls (n = 3-6, Ordinary one-way ANOVA, ** p = < 0.01, *** p = < 0.001).

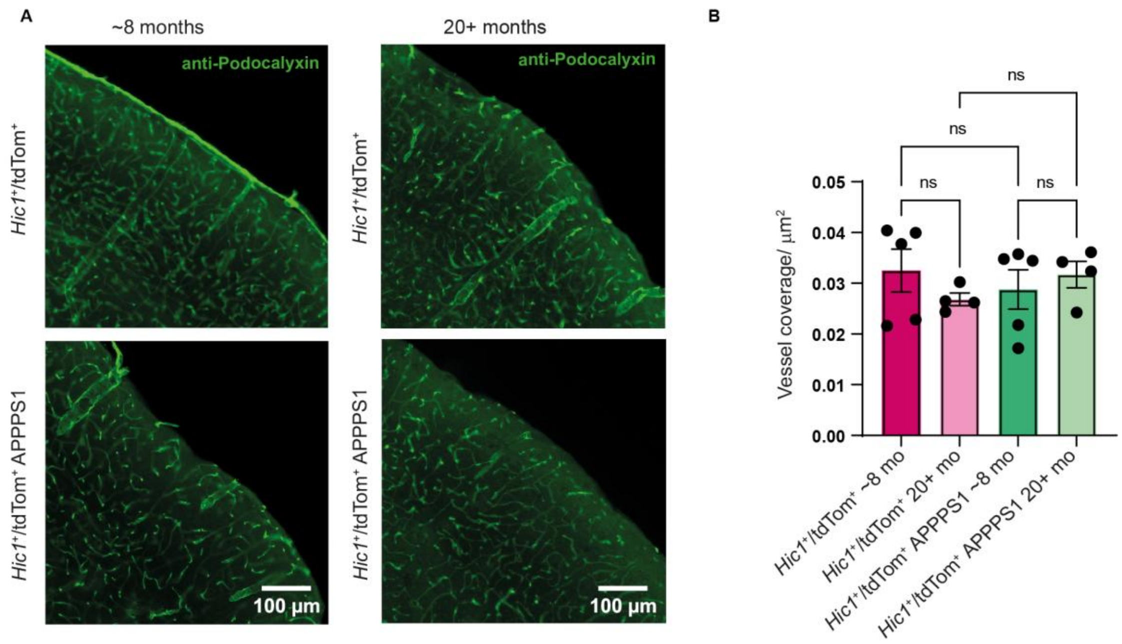


**Extended Data Figure 2. SPCs loss is APPPS1 mice is not due to vessel rarefication.** (A) Representative images of cortex with podocalyxin (green). (B) By analyzing the *Hic1*-tdTomato$^{+}$ somata distribution per vessel length, we found a significant reduction at 20+ months, suggesting that age plus amyloidosis, could lead to a generalized SPC coverage loss (8 months *Hic1*-tdTomato$^{+}$: n=5 mice, 8 months *Hic1*-tdTomato$^{+}$ APPPS1: n=5 mice; 20+ months *Hic1*-tdTomato$^{+}$ n=4 mice; 20+ months *Hic1*-tdTomato$^{+}$ APPPS1: n=4 mice. Ordinary one-way ANOVA Šidak's multiple comparisons).

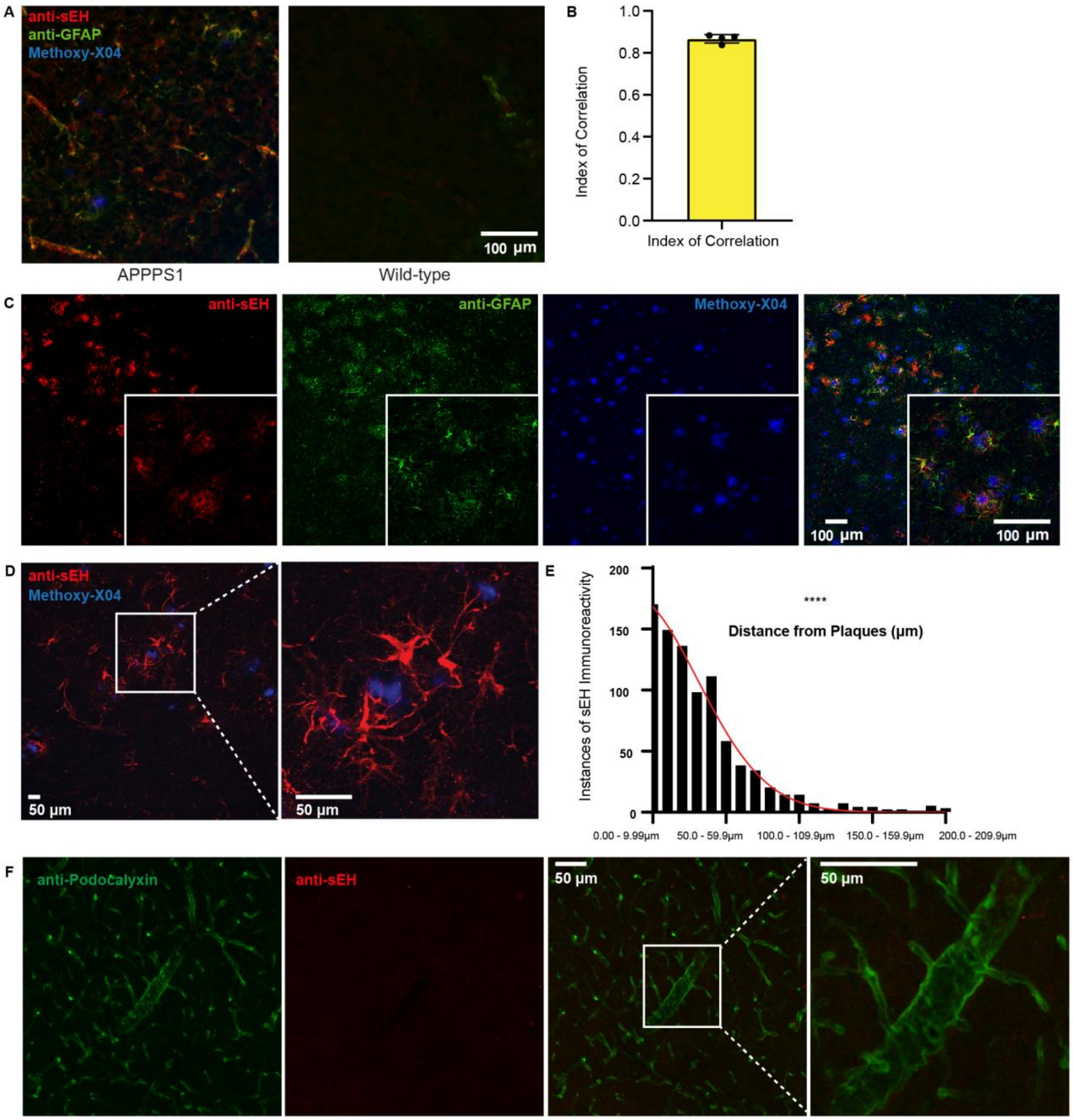


**Extended Data Figure 3. Amyloid pathology drives robust sEH expression in cortical astrocytes.** (A) Representative images of sEH (red) and glial fibrillary acidic protein (GFAP) (green) around plaques (Methoxy-X04, blue) immunoreactivity in APPPS1 cortex compared to wild-type cortex. (B-C) sEH (red) localizes within reactive astrocytes using an antibody against GFAP (green) (n= 4, standard deviation of the index of correlation). (D-E) Frequency distribution of sEH immunoreactivity in proximity to amyloid plaques (Methoxy-X04, blue) reveals a positive correlation between sEH immunoreactivity and amyloid plaques (n = 4, Shapiro–Wilk test, **** p

= <0.0001). (F) Representative image of sEH immunoreactivity (red) on larger vessels (visualized with anti-podocalyxin, green) in wild-type mice.

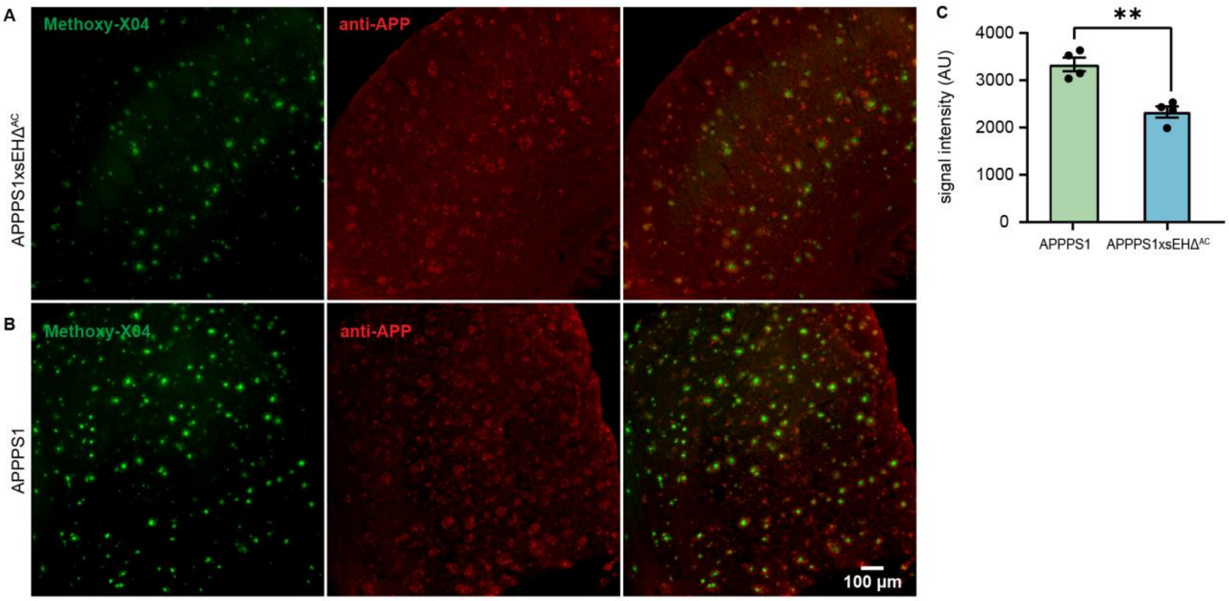


**Extended Data Figure 4. Astrocytic sEH deletion in APPPS1 mice has a neuroprotective effect.** (A-C) APP immunoreactivity (red) was reduced in APPPS1xsEHΔ$^{AC}$ mice (n = 4 two-tailed unpaired t-test, ** p = < 0.01).

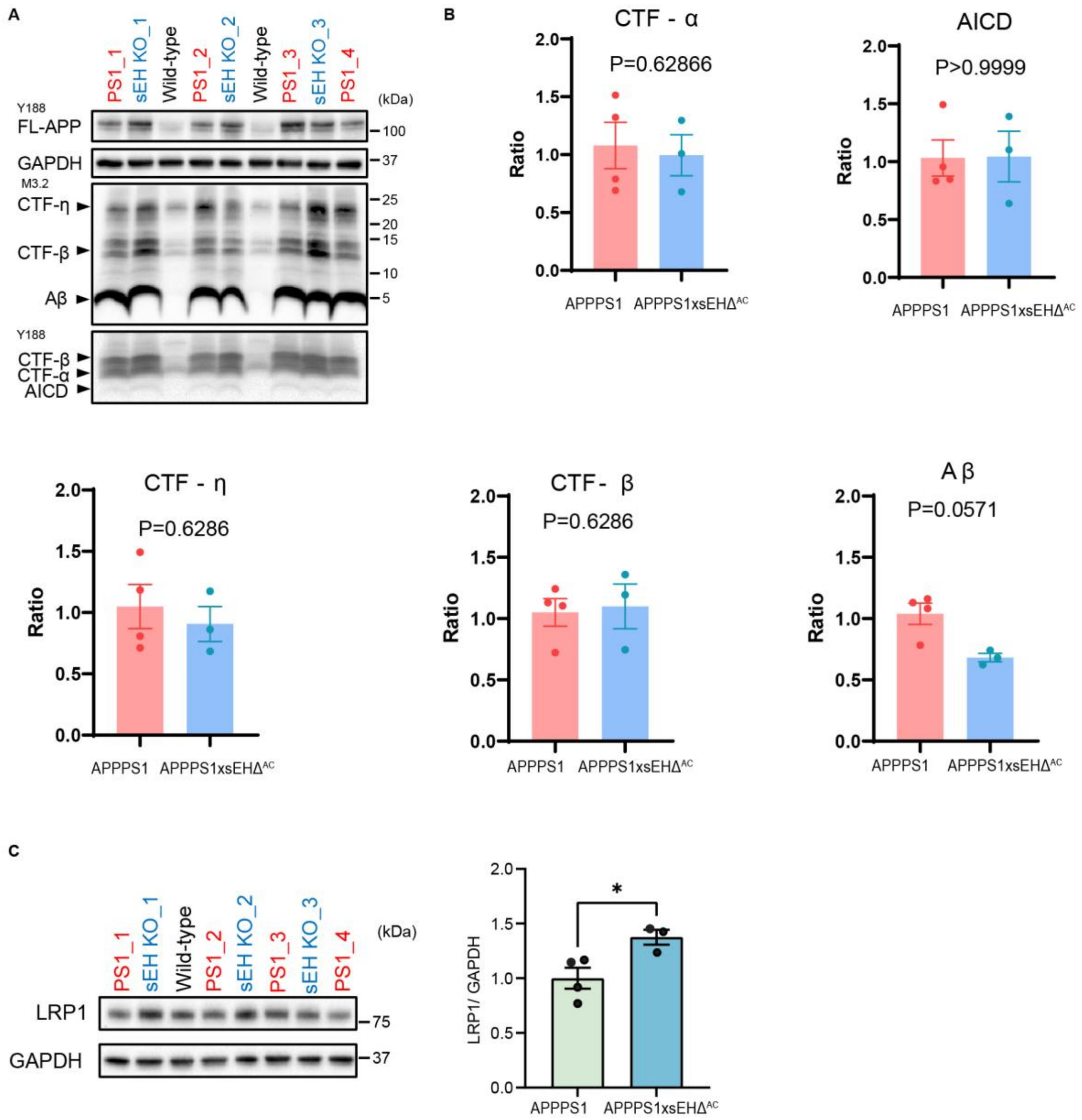


**Extended Data Figure 5. LRP1 contributes to amyloid reduction.** (A-B) APP processing was examined, but shown to be insignificant (n = 3, Mann-Whitney test). (C) LRP1 protein levels are elevated in whole brain homogenates of APPPS1xsEH$\Delta^{AC}$ mice compared to APPPS1 mice (n = 3, two-tailed unpaired t-test, * p = <0.05).

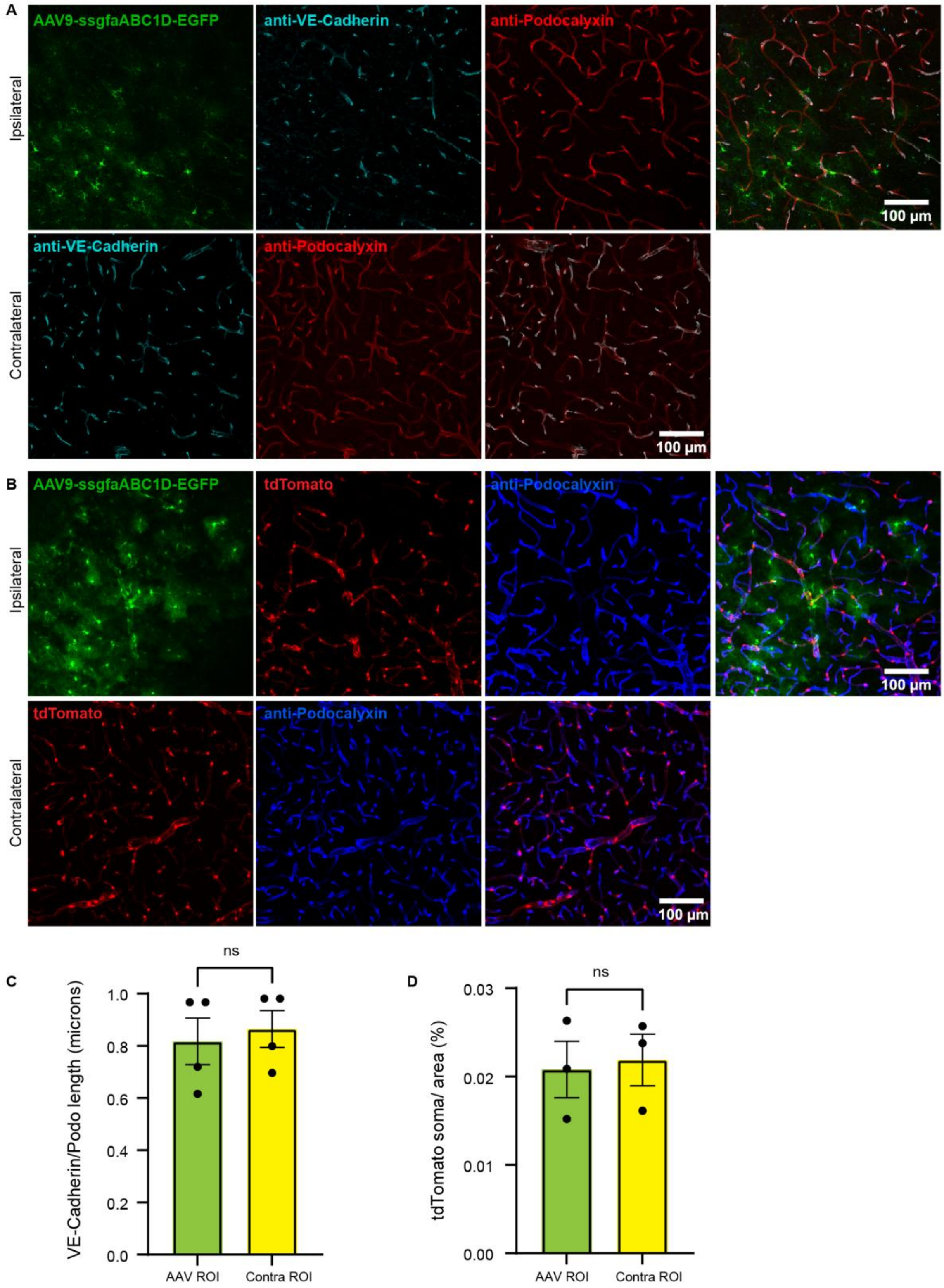


**Extended Data Figure 6. Blood-brain barrier damage driven by viral overexpression of sEH was not seen in sham trials.** (A, C) Control virus, AAV9-ssgfaABC1D-EGFP (green) was injected intracranial, no decrease in relative VE-cadherin (cyan) concentration was found around the viral target region (n = 4, one-tailed paired t-test). (B, D) Similarly *Hic1*-tdTomato$^{+}$ (red) loss was not observed in mice intracranial injected with control virus AAV9-ssgfaABC1D-EGFP (green) (n = 3, one-tailed paired t-test).

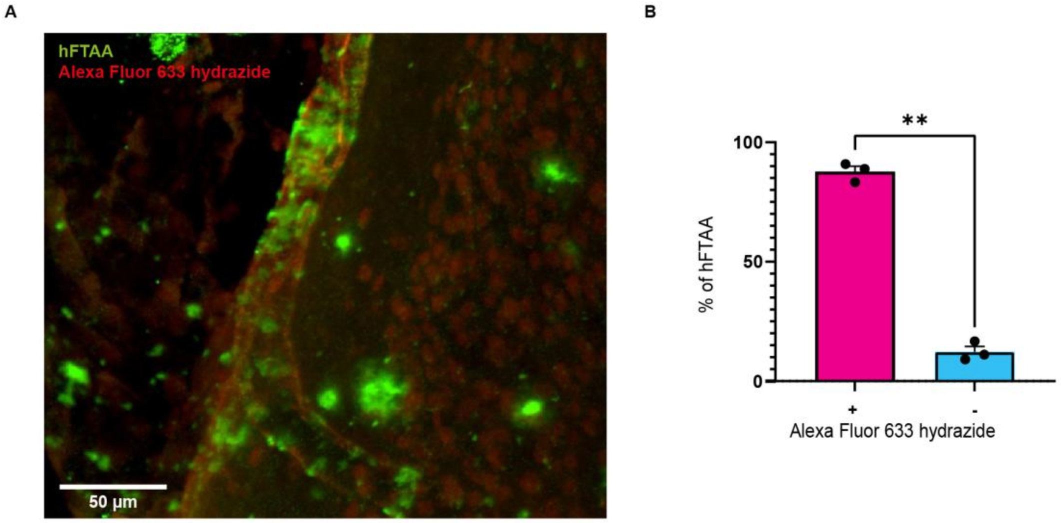


**Extended Data Figure 7. hFTAA is more present in arterioles.** (A,B) Mice were co-injected with Alexa Fluor 633 hydrazide, which selectively binds elastin in arterioles, and hFTAA to visualize CAA. hFTAA was predominantly observed in Alexa Fluor 633 hydrazide-positive arterioles compared to Alexa Fluor 633 hydrazide-negative vessels, indicating preferential amyloid deposition in arterioles (n = 3, two-tailed paired t-test, ** p = < 0.01).
